# A magnesium transporter CorA of *Mycobacterium smegmatis* enhances the tolerance of structurally unrelated antibiotics in the host cells

**DOI:** 10.1101/2024.07.09.602764

**Authors:** Debasmita Chatterjee, A R Daya Manasi, Sumit Kumar Rastogi, Aditya Prasad Panda, Bayomi Biju, Debleena Bhattacharyya, Anindya Sundar Ghosh

**Author notes:** Equal contribution (A R Daya Manasi, Sumit Kumar Rastogi).

## Abstract

Ion transporters or channels are involved in maintaining metal homeostasis in bacterial cells by aiding the movement of metal ions across the cell, which might also facilitate the export of antimicrobials. Ubiquitous magnesium transporter, CorA of *Mycobacterium smegmatis* is well known for its role in maintaining magnesium homeostasis. However, little is known about its involvement in exerting antimicrobial resistance. Here, with the help of molecular genetics, *in vivo* and *in silico* studies we tried to envisage the role of CorA of *M. smegmatis* in antimicrobial resistance of *M. smegmatis* and *E. coli*. Expression of *corA* in *M. smegmatis* and *E. coli* increased the tolerance of the host cells towards various structurally unrelated antibiotics and anti-tubercular drugs. In addition, a significantly lower accumulation of norfloxacin and ofloxacin by the host cells expressing *corA* further indicated its role in enhancing the efflux pump activity. Moreover, the presence of a sub-inhibitory concentration of Mg^2+^ resulted in increased low-level tolerance towards the tested drugs. Furthermore, CorA enhanced the biofilm-forming ability of cells expressing it. Overall, we speculate that magnesium transporter CorA facilitates multi-drug efflux activity of the host cells where Mg^2+^ might act as a facilitator in the process.

**IMPORTANCE:** Magnesium acts as a co-factor for various biochemical and physiological reactions, such as protein synthesis, cell membrane integrity, nucleic acid synthesis, etc. Metal transporters maintain metal homeostasis by regulating the uptake, efflux, or transportation of metals in certain necessary cellular compartments. In bacteria, magnesium ion (Mg^2+^) is mainly supplied by the CorA protein which is a ubiquitous family of transport proteins and extensively studied in *E. coli* and *Salmonella sp*. However, little is known about the functional relationship of metal transporters of *Mycobacterium sp* with extrusion of antibiotics, and their involvement in stress tolerance. Here, we report CorA (MSMEG_5056), a magnesium transporter of *Mycobacterium smegmatis* in influencing the extrusion of multiple structurally unrelated classes of drugs and enhancing the biofilm formation of *E. coli* and *Mycobacterium smegmatis*.

Graphical Abstract:
Hypothetical model of Antibiotic export by CorA.
Antibiotics bind to the closed state of the protein (left). During the transition to the open state (right), the to-and-fro motion between multiple open states drives the efflux of the antibiotic while facilitating the import of Mg^2+^. The colors of the models correspond to the chain ID. The bottom views of both the closed and open states are shown in the rectangular box, with the color indicated by their respective chain IDs.

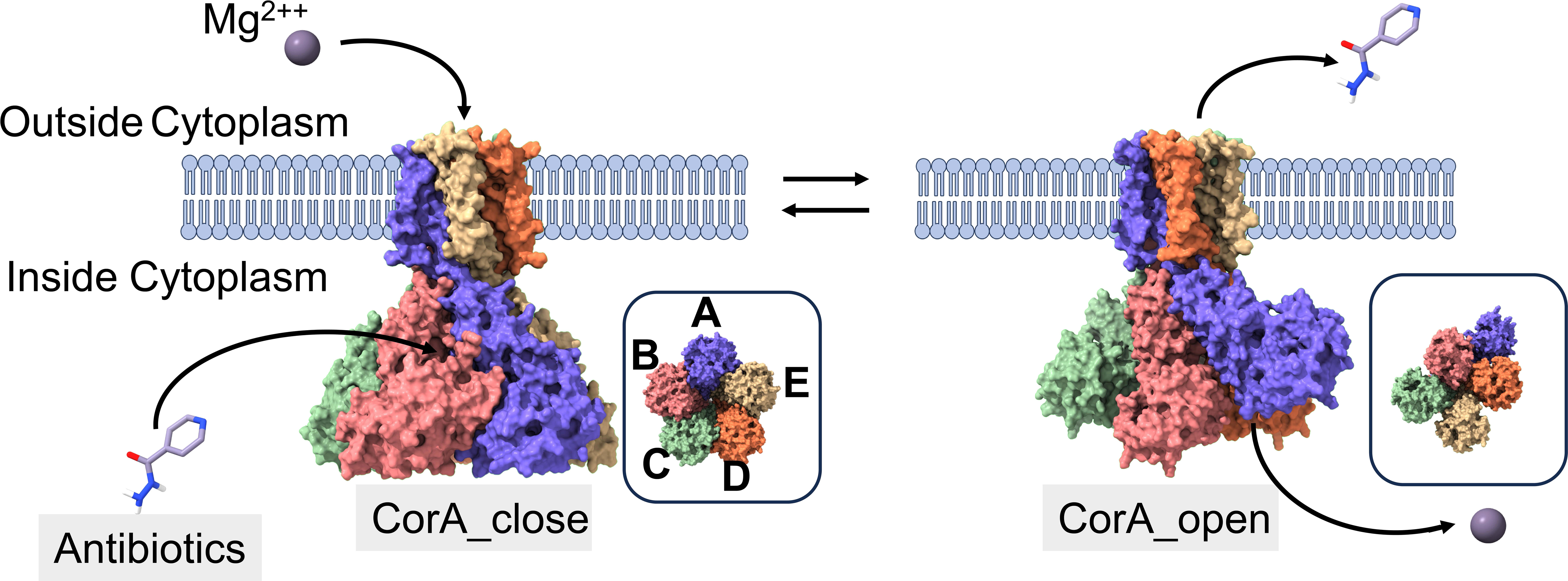

## INTRODUCTION

Despite recent advancements in the epidemiology of tuberculosis (TB) and knowledge of its causative organism *Mycobacterium tuberculosis* (Mtb), TB remains one of the deadliest diseases caused by a single infectious agent. The emergence of MDR-TB and XDR-TB poses a major health threat worldwide (1). Drug resistance in Mtb is primarily attributed to the presence of thick lipid-rich membranes, active efflux systems, chromosomal mutations affecting enzymatic performances, and drug target modifications (2, 3).

So far, the metal transporters of *Mycobacterium sp* have not been studied extensively. Metallo-enzymes require co-factors like iron, copper, and zinc which play key structural and catalytic roles in the cells (4). Magnesium is required for macromolecule synthesis and protein synthesis (Takezawa et al., 2016; Yang et al., 2004). Magnesium and iron are also required for the survival of the pathogen within macrophages (7). To maintain the cellular homeostasis of magnesium, cells require a transport system. There are three classes of known transport systems, namely, CorA, MgtA/B, and MgtE, which are involved in the transportation of magnesium ions in and out of the cells (8). In eubacteria and archaea, mostly CorA and MgtE are present, whereas MgtA/B is present in those cells that are devoid of the former two proteins (8). Like MgtA and MgtB, CorA also follows the electrochemical gradient across the cytoplasmic membrane to transport its substrates (Niegowski, Eshaghi 2007). CorA mediates the transport of Mg^2+,^ Co^2+^, and Ni^2+^ (9). In *Salmonella typhimurium*, mutation of CorA led to the attenuation of virulence even though it has two additional magnesium transporters (10).

It is reported that metals may also play a key role in antibiotic resistance via the development of cross-resistance (11). In *Pseudomonas aeruginosa,* heavy metal efflux pump CzcCBA imparts cross-resistance to Carbapenem (12). In *Listeria monocytogenes* efflux pump can throw out zinc, cobalt, and cadmium as well as antibiotics like erythromycin and clindamycin (13). A genome-wide study in *Acinetobacter baumannii* showed that efflux pumps are involved in the resistance to both, heavy metals and antibiotics (14). On the other hand, the roles of transmembrane ion transporters and their involvement in providing antimicrobial resistance are relatively less known in mycobacterial species.

Metal homeostasis has been co-related with combating antibiotic stress (15). The higher magnesium flux influences the growth and survival of bacteria that are exposed to antibiotics targeting the ribosome (16). Ribosomes comprise numerous rRNA units where Mg^2+^ ions stabilize the ribosomal complex (Akanuma *et al.* 2018). Magnesium plays an important role in combating the ribosome-inhibiting antibiotic stress (16). Therefore, combining antibiotics with the inhibitors that are targeting these transporters may enhance the efficacy of the antibiotics.

In the present study, with the help of molecular and *in vivo* expression analyses such as antibiotic sensitivity assessments, intracellular dye, and antibiotic accumulation, semi-quantitative biofilm formation and growth curve, and *in silico* analyses (e.g. molecular docking), we attempted to investigate the role of CorA of *Mycobacterium smegmatis*, a magnesium transporter, in providing antibiotic tolerance as well as affecting biofilm formation in *Mycobacterium smegmatis* and *E. coli*.

## RESULTS

### CorA influences the resistance of *E. coli* cells toward structurally unrelated classes of antibiotics

To ascertain the role of CorA in drug resistance, the drug susceptibility profile of the cells expressing *corA* was determined against various classes of antibiotics. CorA of *M. smegmatis* was expressed in *E. coli* with 0.1% arabinose (**Figure S2A**). The over-expression of *corA* in *E. coli* cells decreased the susceptibility of the host cells towards multiple antibiotics namely, norfloxacin (4 fold), ofloxacin (4 fold), sparfloxacin (2 fold), ciprofloxacin (4 fold), amikacin (4 fold), gentamicin (4 fold), apramycin (2 fold) (**Table 1**). According to the literature, CorA is a known magnesium transporter (30). In the presence of a sub-inhibitory concentration of magnesium (100ppm) (MgCl_2_), the susceptibilities of the host cells were decreased by 4 – 8 fold towards all the tested antibiotics (**Table 1**). Further, to ascertain that the change in susceptibility was due to magnesium, the assay was performed in the presence of zinc chloride (ZnCl_2_). However, the susceptibility values were the same as the values determined without the presence of any other compound, nullifying the effect of chloride ions in the experimental setup. Interestingly, after the addition of CCCP, a potent uncoupler of proton motive force (pmf), the sensitivity of antibiotics like norfloxacin, ofloxacin, ciprofloxacin, and gentamicin was increased by 2 fold whereas sensitivity remained unaltered for sparfloxacin, amikacin and apramycin in both the set of cells whether expressing *corA* or deficient of *corA* (**Table 2**). As a whole, the decrease in susceptibility of the host *E. coli* cells towards different classes of antibiotics indicates the involvement of CorA in enhancing multi-drug resistance where CCCP might have an inhibitory effect.

**Table 1:**
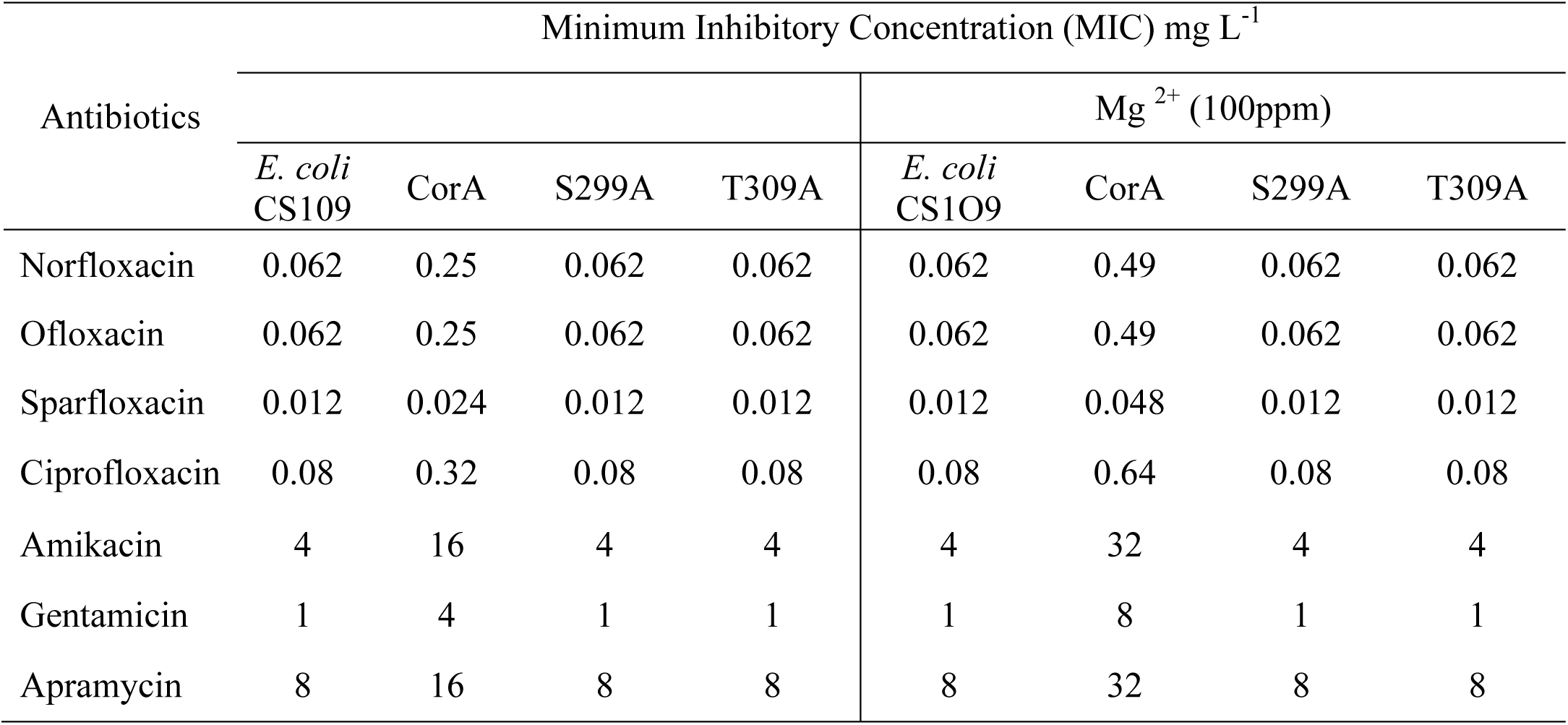
Minimum Inhibitory Concentrations of various unrelated classes of antibiotics against CS109 cells carrying pBAD18 Cam vector (control of *E. coli* cells), and CS109 cells harboring CorA (pDCorA) and its mutants in the presence and absence of Mg^2+^.

**Table 2:**
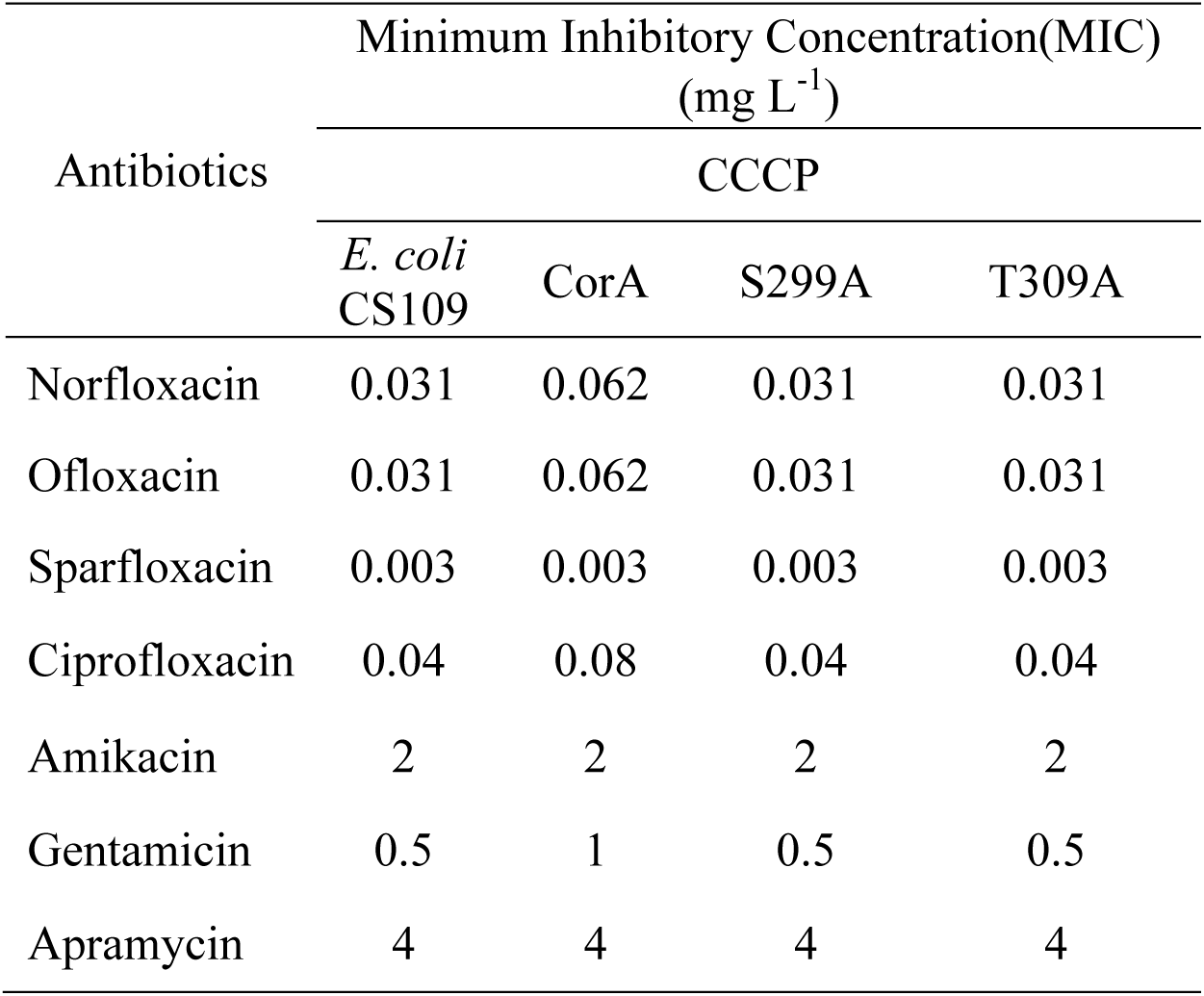
Minimum Inhibitory Concentrations of various unrelated classes of antibiotics against CS109 cells carrying pBAD18 Cam vector (control of *E. coli* cells), and CS109 cells harboring CorA (pDCorA) and its mutants in the presence of CCCP.

### *corA* expression in *E. coli* cells reduces norfloxacin accumulation

To assess whether the increase in tolerance of the host cells towards different antibiotics was due to the enhancement of efflux pump activity, the time-dependent intracellular drug accumulation assay of norfloxacin was performed. It was noted that the accumulation of norfloxacin by *E. coli* cells over-expressing *corA* was lower than that of the cells carrying empty pBAD-18Cam. It was observed that within 15 min exposure to norfloxacin, there was a relative decrease in antibiotic accumulation by 44% in the cells over-expressing *corA*, than that of the control. However, the accumulation of norfloxacin was markedly increased within 15 min upon the addition of CCCP in all the tested strains, as observed due to a rapid increase in fluorescence signal (**Figure 1**). Therefore, we infer that the increase in tolerance of *corA* over-expressing cells towards antibiotics is apparently due to the decrease in the accumulation of antibiotics.

**Figure 1:**
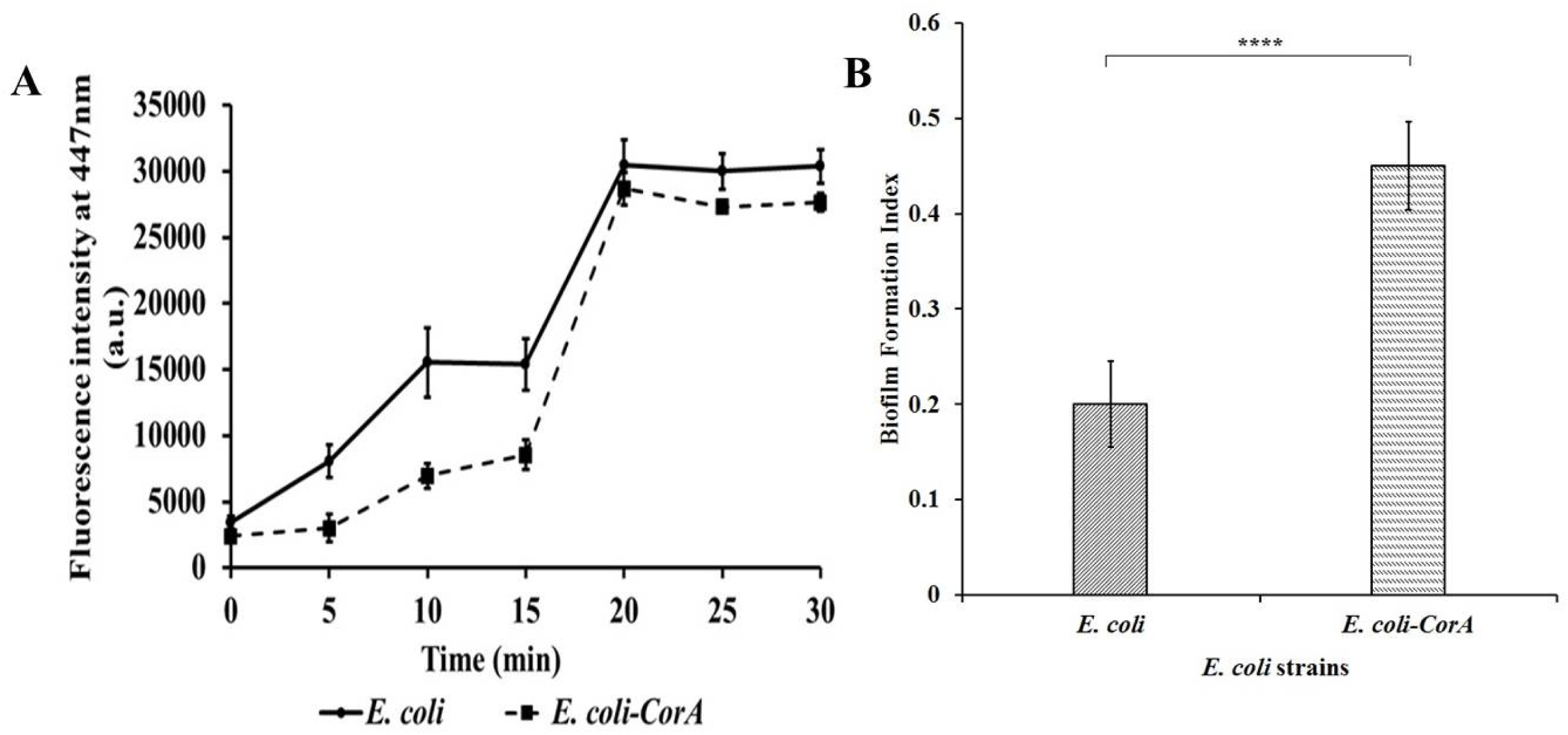
Intracellular accumulation of norfloxacin in *E. coli* cells. (A) harboring CorA in comparison to control cells (harboring empty vector) with respect to time. Norfloxacin was added at 0^th^ min. Sample aliquots were drawn at every 5 min intervals for 30 min and CCCP was added at 15^th^ min. (B) Relative efflux was calculated after exposing the *E. coli* cells to norfloxacin for 10 min. Two-tailed unpaired Student’s *t*-test was performed with the control and test datasets where ***** denotes P<0.0003. The error bars indicate mean±SD.

### Deletion of *corA* hampers growth of *M. smegmatis*

CorA was deleted from the *M. smegmatis* cells (**Figure S1**) following the homologous recombination method. The deletion was confirmed by selecting the positive colonies on 7H11-hygromycin plates supplemented with 10% sucrose. The transformants obtained were named *ΔmsCorA*. Further deletion confirmation was done by amplifying the genomic DNA of CorA-deleted *M. smegmatis* with a combination of multiple primers (**Figure S1 and Table S1**). Thereafter CorA cloned in pMIND shuttle vector was electroporated in *M. smegmatis* CorA deleted cells. The transformants were *ΔmsCorA::CorA*. The growth pattern was analyzed by comparing the same among the deletion mutant, CorA-complemented, and wild-type strains. We observed a ∼19% increase in generation time and a 13% lower growth rate in CorA deleted strain (*ΔmsCorA*) (3.6 h and 0.20 h^-1^) as compared to the parent strain (wild-type *M. smegmatis*) and CorA complemented strain (*ΔmsCorA::CorA*) (3.03h and 0.23 h^-1^) (**Figure 2**). The results indicate that the deletion of CorA affects the growth of the bacteria.

**Figure 2:**
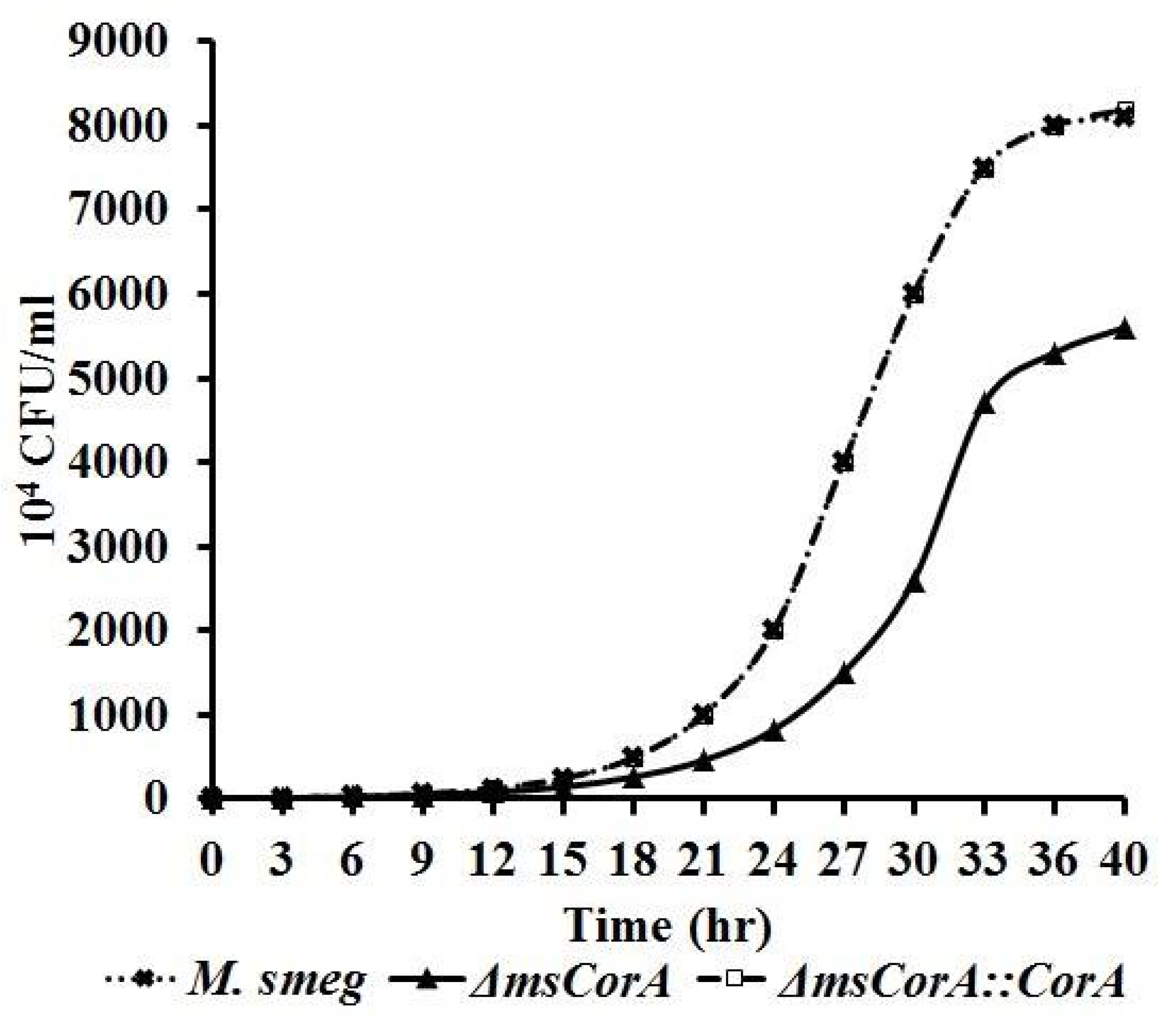
Growth analysis of *M. smegmatis* and CorA knock-out *M. smegmatis* strains: The strains were inoculated (McFarland number 0.5) into Middlebrook 7H9 medium (supplemented with OADC enrichment and appropriate antibiotics), and the CFU per milliliter were counted every 3 h for 40 h. The growth rate and doubling time were calculated for each replicate using the linear part of the log phase.

### Expression of *corA* increases antibiotic tolerance of *M. smegmatis* cells

As we observed that over-expression of *corA* increased the tolerance of the *E. coli* host towards structurally unrelated classes of antibiotics and decreased the accumulation of norfloxacin, we were interested to see the effect of *corA* expression in its native host *M. smegmatis* and accordingly, we proceeded. It was observed that the tolerance of *ΔmsCorA* towards different antibiotics such as norfloxacin (4 fold), ofloxacin (4 fold), sparfloxacin (2 fold), ciprofloxacin (4 fold), amikacin (4 fold), gentamicin (4 fold), apramycin (2 fold), isoniazid (4 fold) and rifampicin (8 fold) was reduced (**Table 3**). Interestingly, the susceptibility levels were restored when the CorA-deleted mutant was complemented with CorA cloned in the pMIND vector and expressed with 20 ng/µL tetracycline (**Figure S2B**). Since, CorA is an established magnesium transporter in *M. smegmatis*, the effect of magnesium on the tolerance levels of the cells expressing *corA* towards antibiotics was studied by supplementing 100 ppm of magnesium into the experimental setup. It was observed that in the presence of sub-inhibitory concentration of magnesium, the susceptibilities of the host cells expressing *corA* further decreased by 4 – 16 fold (**Table 3**). After addition of CCCP the sensitivities of antibiotics like norfloxacin, ofloxacin, ciprofloxacin, and gentamicin were increased by 2 fold, though the sensitivities did not change for sparfloxacin, amikacin, isoniazid, rifampicin and apramycin in the cells expressing *corA* (*ΔmsCorA::CorA* and wild type *M. smegmatis*) and cells deficient of CorA (*ΔmsCorA*) (**Table 4**). Therefore, the decrease in susceptibility levels of the host cells expressing *corA* (*ΔmsCorA::CorA* and wild-type *M. smegmatis*) implies the involvement of CorA in the antimicrobial resistance of *M. smegmatis*.

**Table 3:**
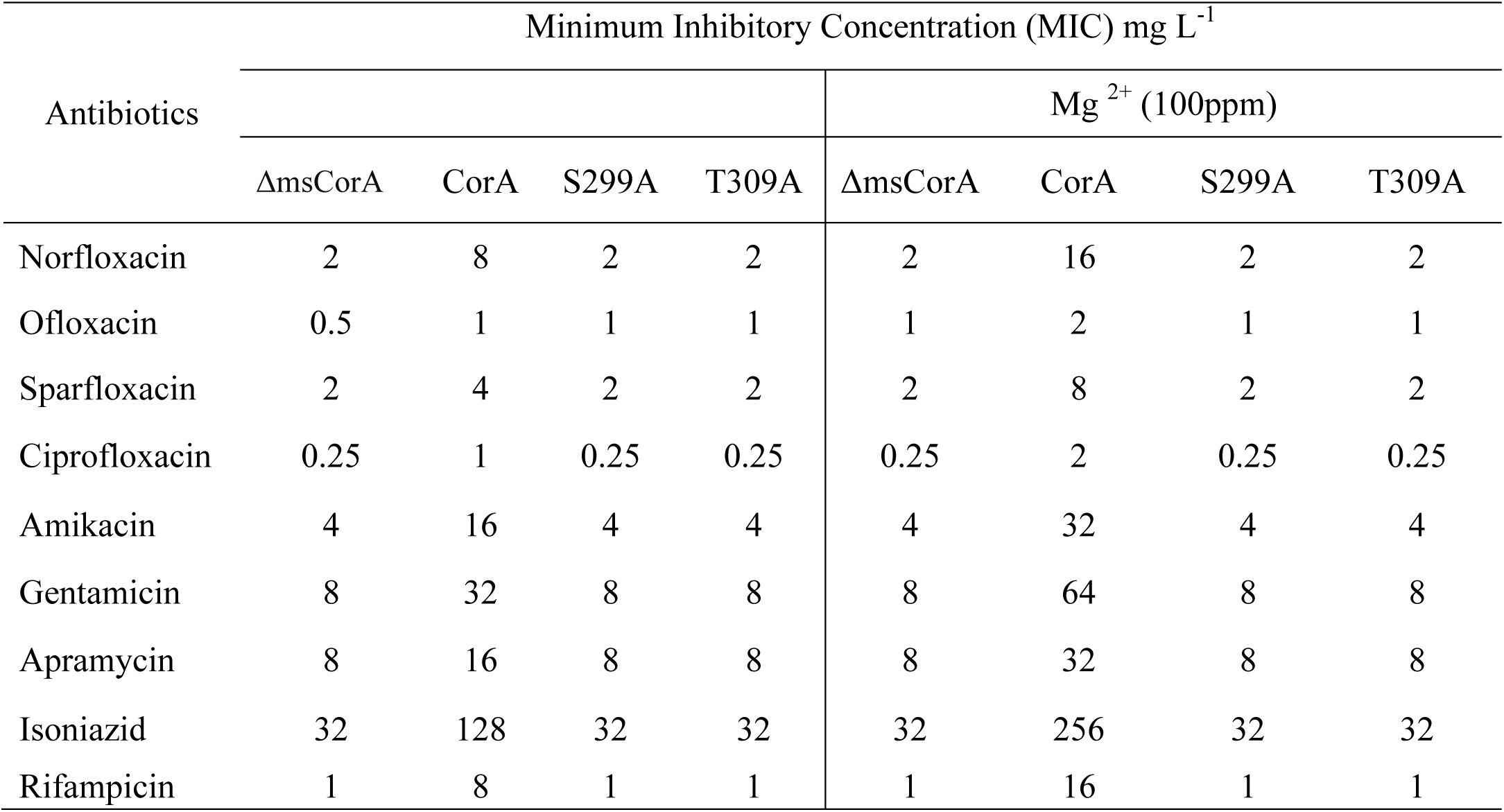
Minimum Inhibitory Concentrations of various unrelated classes of antibiotics against, *M. smegmatis* cells harbouring pMIND vector (control of *M. smegmatis* cells) and CorA (pMCorA) and its mutants in the presence and absence of Mg^2+^.

**Table 4:**
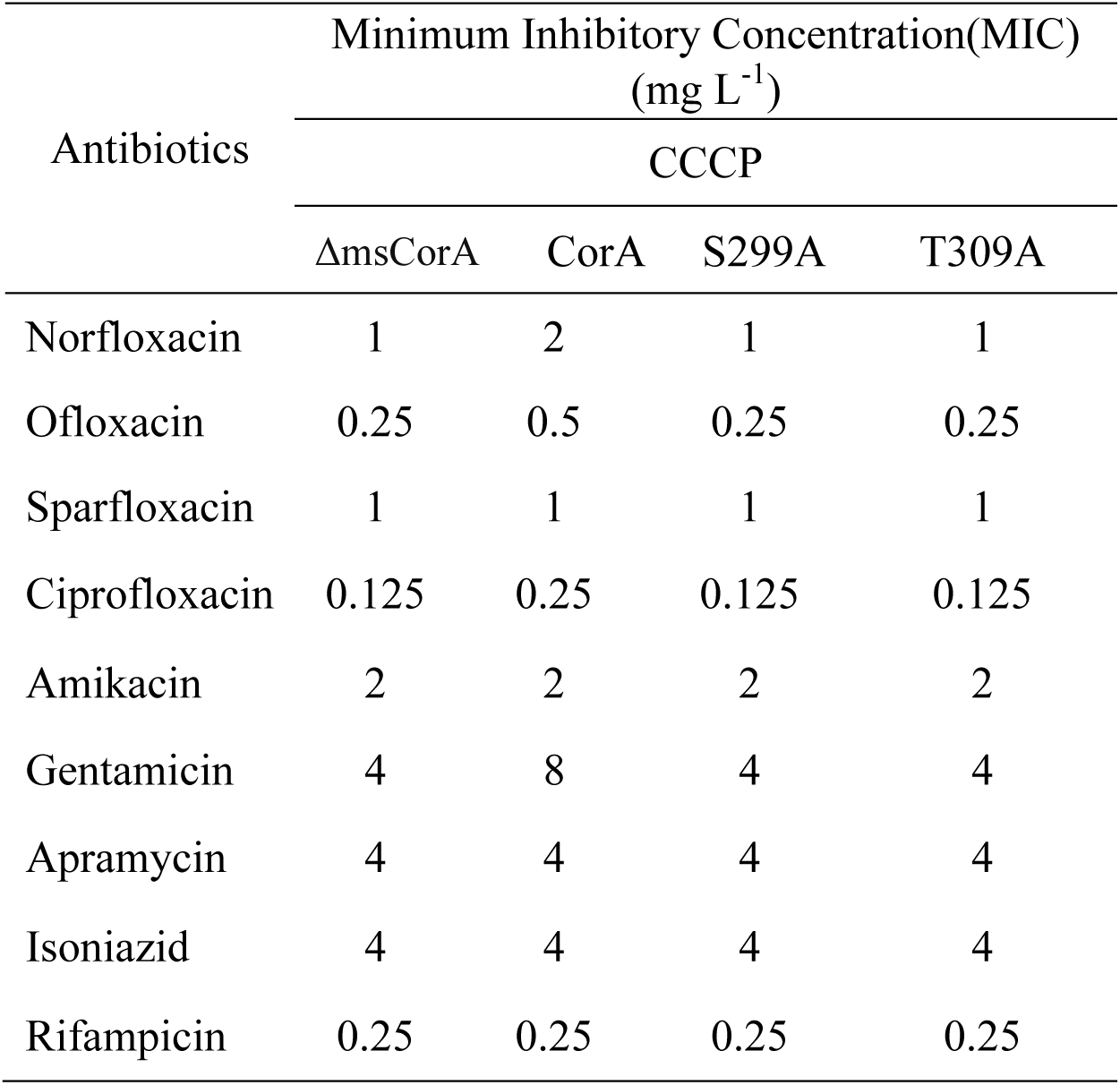
Minimum Inhibitory Concentrations of various unrelated classes of antibiotics against, *M. smegmatis* cells harbouring pMIND vector (control of *M. smegmatis* cells) and CorA (pMCorA) and its mutants in the presence of CCCP.

### Intracellular accumulations of antibiotics/dye are reduced upon expression of *corA* in *M. smegmatis*

To denote that the increase in tolerance of the host cells towards antibiotics was due to the influence of CorA on the efflux activity of *M. smegmatis*, we performed an intracellular antibiotics/dye accumulation assay. It was observed that host cells harboring CorA (*ΔmsCorA::CorA* and wild type *M. smegmatis*) accumulated relatively lesser EtBr, ofloxacin, and norfloxacin as compared to CorA deleted strain (*ΔmsCorA*). It was noted that within 8 min of exposure of EtBr to CorA cells, the accumulation levels of EtBr were reduced by 40% (**Figure 3**). Further, the accumulation levels of norfloxacin and ofloxacin by CorA complemented mutant and wild-type *M. smegmatis* cells relatively decreased by 48% - 50% within the first 10 min than in the CorA deletion mutant (**Figure 4A and 4C, 5A, and 5C**). It was previously demonstrated that the tolerance of the host cells expressing *corA* was increased at a low level towards the tested antibiotics in the presence of magnesium. Therefore, the influence of magnesium on the accumulation levels of the antibiotics was determined for the host cells (30). In the presence of sub-inhibitory concentration of magnesium, the accumulation levels of norfloxacin and ofloxacin by the host cells expressing c*orA* was reduced by more than 50% as compared to the cells lacking CorA (**Figure 4B and 4D, 5B and 5D**). There was a consistent and significant decrease in the accumulation of antibiotics in the cells harboring CorA as observed from the relative efflux data that indicates the role of CorA in influencing the multi-drug efflux activity.

**Figure 3:**
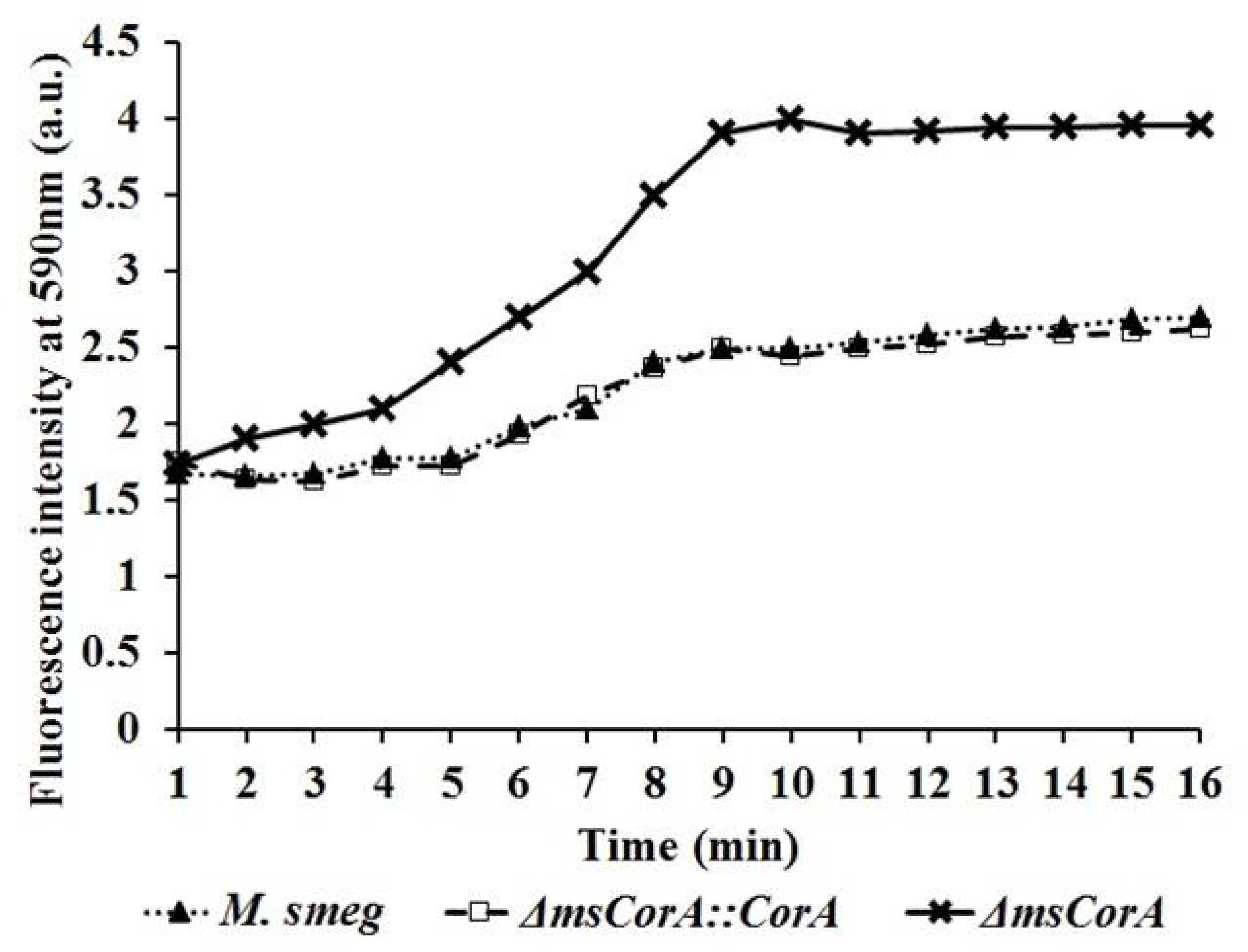
Intracellular accumulation of EtBr in *M. smegmatis* cells: EtBr accumulations in *M. smegmatis* cells were determined for 16 min. The substrate was added to the cells at the 0^th^ min. Fluorescence intensity at 590 nm was determined every 1 min for 16 min.

**Figure 4:**
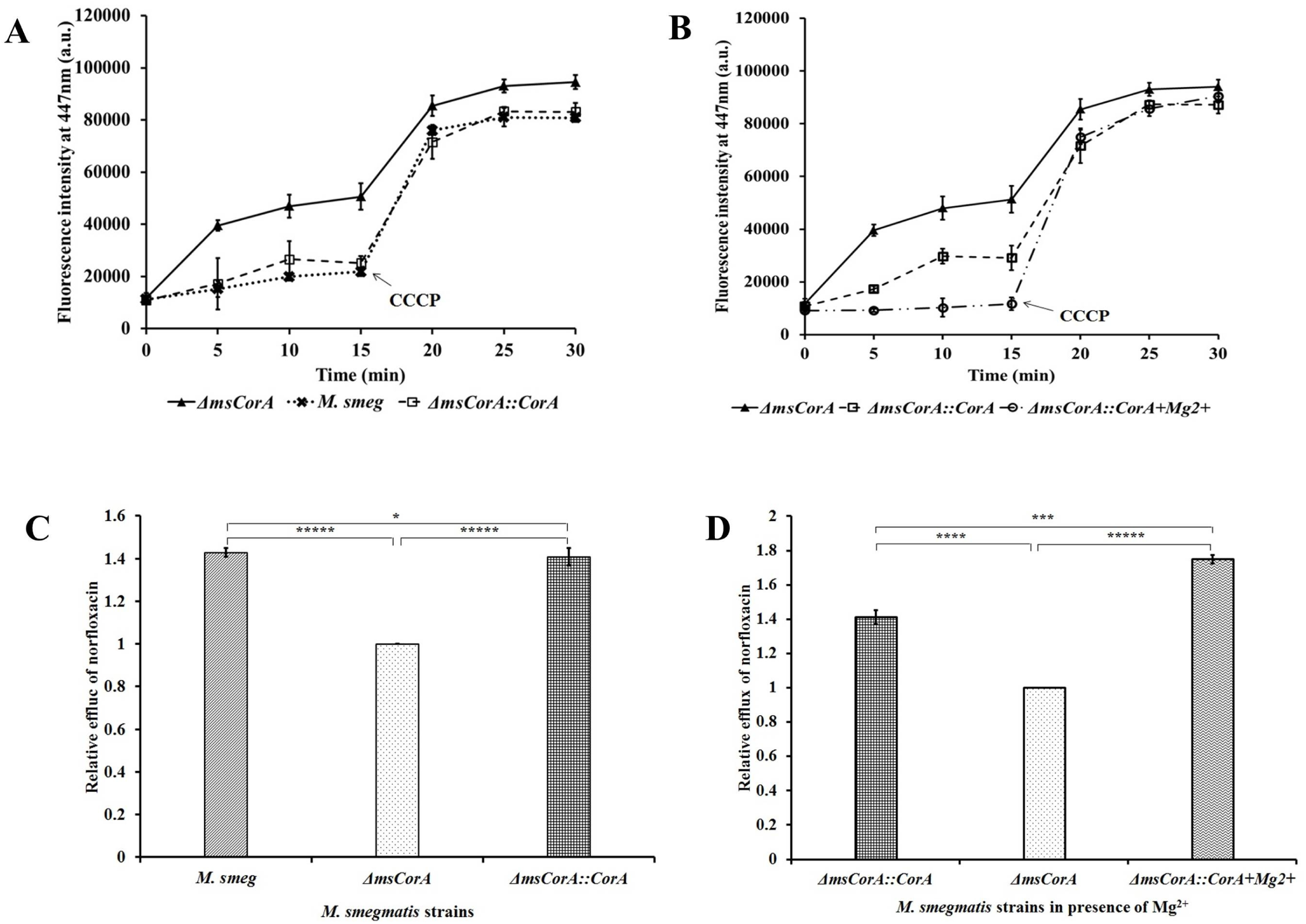
**Intracellular norfloxacin accumulation** in *M .smegmatis* cells harboring CorA and devoid of CorA with respect to time. (A) Norfloxacin was added at 0^th^ min. Sample aliquots were drawn at every 5 min interval for 30 min and CCCP was added at 15^th^ min. (B) Norfloxacin accumulation in the presence of sub-inhibitory concentration of Mg^2+^ by host cells was also determined. (C) and (D) Relative efflux was calculated after exposing the *M. smegmatis* cells to norfloxacin both in the absence and presence of Mg^2+^ for 10 min. Two-tailed unpaired Student’s *t*-test was performed with the control and test datasets where* denotes P<3, *** denotes P<0.03, **** denotes P<0.003 and ***** denotes P<0.0003. The error bars indicate mean±SD

Further, it was noticed that after the addition of CCCP, the accumulation levels in the experimental cells as well as the control cells rapidly increased (**Figures 4 and 5**). Also, the differences in the levels of accumulation of antibiotics were relatively decreased by all the tested cells. Therefore, we hypothesize that likewise in *E. coli*, CorA of *M. smegmatis* influences the efflux activity of the host cells where CCCP might act as an inhibitor.

**Figure 5:**
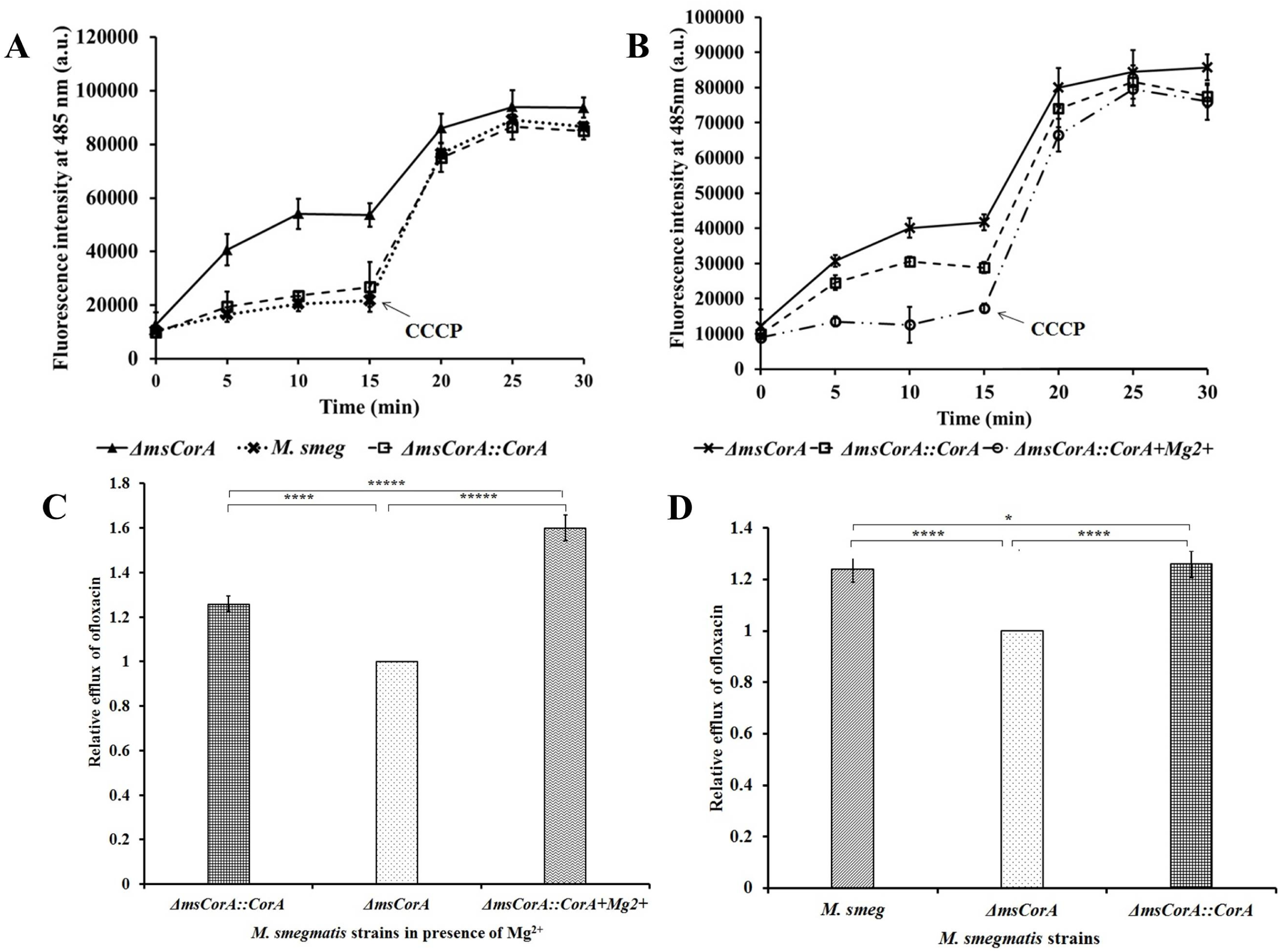
**Intracellular ofloxacin accumulation** in *M .smegmatis* cells harboring CorA and devoid of CorA with respect to time. (A) Ofloxacin was added at 0^th^ min. Sample aliquots were drawn at every 5 min intervals for 30 min and CCCP was added at 15^th^ min. (B) Ofloxacin accumulation in the presence of sub-inhibitory concentration of Mg^2+^ by host cells was also determined. (C) and (D) Relative efflux was calculated after exposing the *M. smegmatis* cells to ofloxacin both in the absence and presence of Mg^2+^ for 10 min. Two-tailed unpaired Student’s *t*-test was performed with the control and test datasets where* denotes P<3, *** denotes P<0.03, **** denotes P<0.003 and ***** denotes P<0.0003. The error bars indicate mean±SD.

#### The transporter function of CorA is affected by the substitution mutations S299A and T309A

Mary Ann Szegedy determined that the hydroxyl-bearing residues S260, and T270 were important for the metal transport through *S. typhimurium* CorA (31). S260 and T270 mutants in *S. typhimurium* had no measurable cation transport. Therefore, to study the effect of these conserved hydroxyl residues of *M. smegmatis* CorA on the antimicrobial efflux activity; site-directed mutagenesis of hydroxyl conserved residues S299 and T309 was performed. The single amino acid substitution of S299A and T309A resulted in the loss of antimicrobial efflux activity of CorA. The mutations led to a complete loss of resistance of the host cells expressing *corA* towards fluoroquinolones, aminoglycosides, and anti-tubercular drugs as compared to the wild-type cells (**Tables 1-4**). Resistance of the host cells harboring CorA against all the tested antibiotics in the presence of Mg^2+^ was also completely nullified though there was no change in the growth pattern of the cells expressing mutated *corA*. The loss of drug efflux activity of CorA due to the mutations was also noted in intracellular norfloxacin and ofloxacin accumulation assays (**Figure 6**). The accumulation of the tested antibiotics by the host cells harboring mutated CorA was similar to the cells that were deficient in CorA. Therefore, it was observed that the conserved hydroxyl residue mutants led to attenuation of the transport capability of CorA.

**Figure 6:**
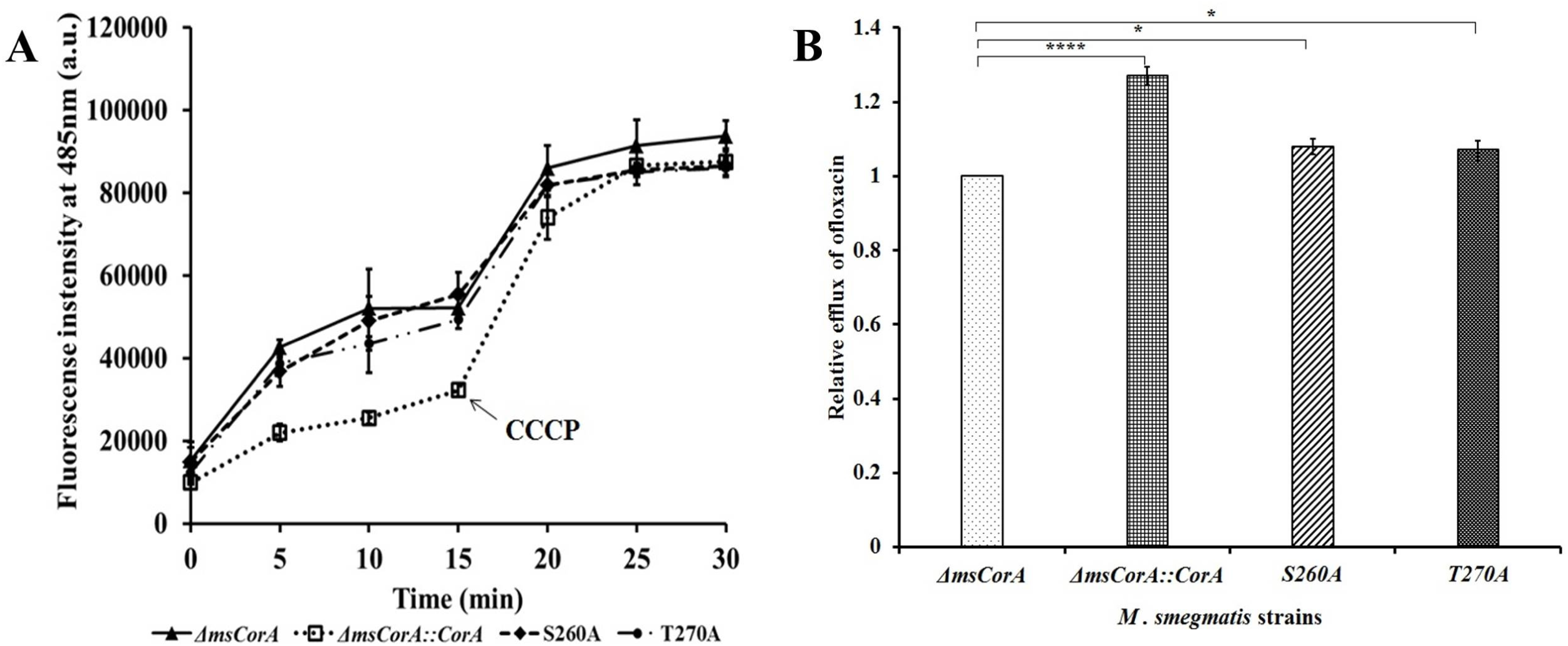
**Intracellular ofloxacin accumulation** in *M .smegmatis* cells harboring CorA and devoid of CorA along with its mutants S299A and T270A with respect to time. (A) Ofloxacin was added at 0^th^ min. Sample aliquots were drawn at every 5 min interval for 30 min and CCCP was added at 15^th^ min. (B) Relative efflux was calculated after exposing the *M. smegmatis* cells to ofloxacin for 10 min. Two-tailed unpaired Student’s *t*-test was performed with the control and test datasets where* denotes P<3 and **** denotes P<0.003. The error bars indicate mean±SD.

### CorA influences the biofilm-forming ability of host cells

It has been demonstrated that efflux pumps play a pivotal role in biofilm formation (32, 33). The pumps are associated with the release of EPS (extra polymeric substances) and quorum sensing molecules, they indirectly regulate the genes for biofilm formation, and efflux of harmful toxins and antimicrobials, thereby promoting aggregation by releasing adhesins (34). To determine the effect of CorA on the biofilm-forming ability of *E. coli* and *M. smegmatis*, a semi-quantitative biofilm quantification assay was performed. It was noted that the Biofilm Formation Index (BFI) for control and *corA* expressing cells for *E. coli* were 0.2 and 0.45, respectively, and for *M. smegmatis* cells the BFIs were 1.19 for wild-type *M. smegmatis* cells, and 1.21 for CorA complemented cells and 0.61 for CorA deleted cells (**Figure 7A and 8A**) (**Figure S3**). Therefore, a relative increase of BFI by 2 fold was observed in the cells expressing *corA*. In the presence of Mg^2+^ the BFI for *corA*-expressing cells increased by 2 to 4 fold for both *E. coli* and *M. smegmatis* cells (**Figure 7B and 8B**). According to the literature, CCCP reduced the biofilm formation of *Acinetobacter baumannii* and *Klebsiella pneumoniae* (35, 36). Likewise in the present study, when 100 µM CCCP was introduced into the experimental setup, the difference in BFI between the control and experimental strain was reduced to less than 2 fold for *E. coli* cells whereas the difference was almost negligible for *M. smegmatis* cells (**Figure 7C and 8C**). It was also observed that the biofilms formed by the wild type (*M. smegmatis*) and CorA complemented cells (*ΔmsCorA::CorA*) were relatively 26% more viable than those of CorA deleted cells (**Figure 9**). This implies that CorA might play a role in influencing the biofilm-forming ability of the microorganisms and CCCP might have an inhibitory effect on the biofilm formation.

**Figure 7:**
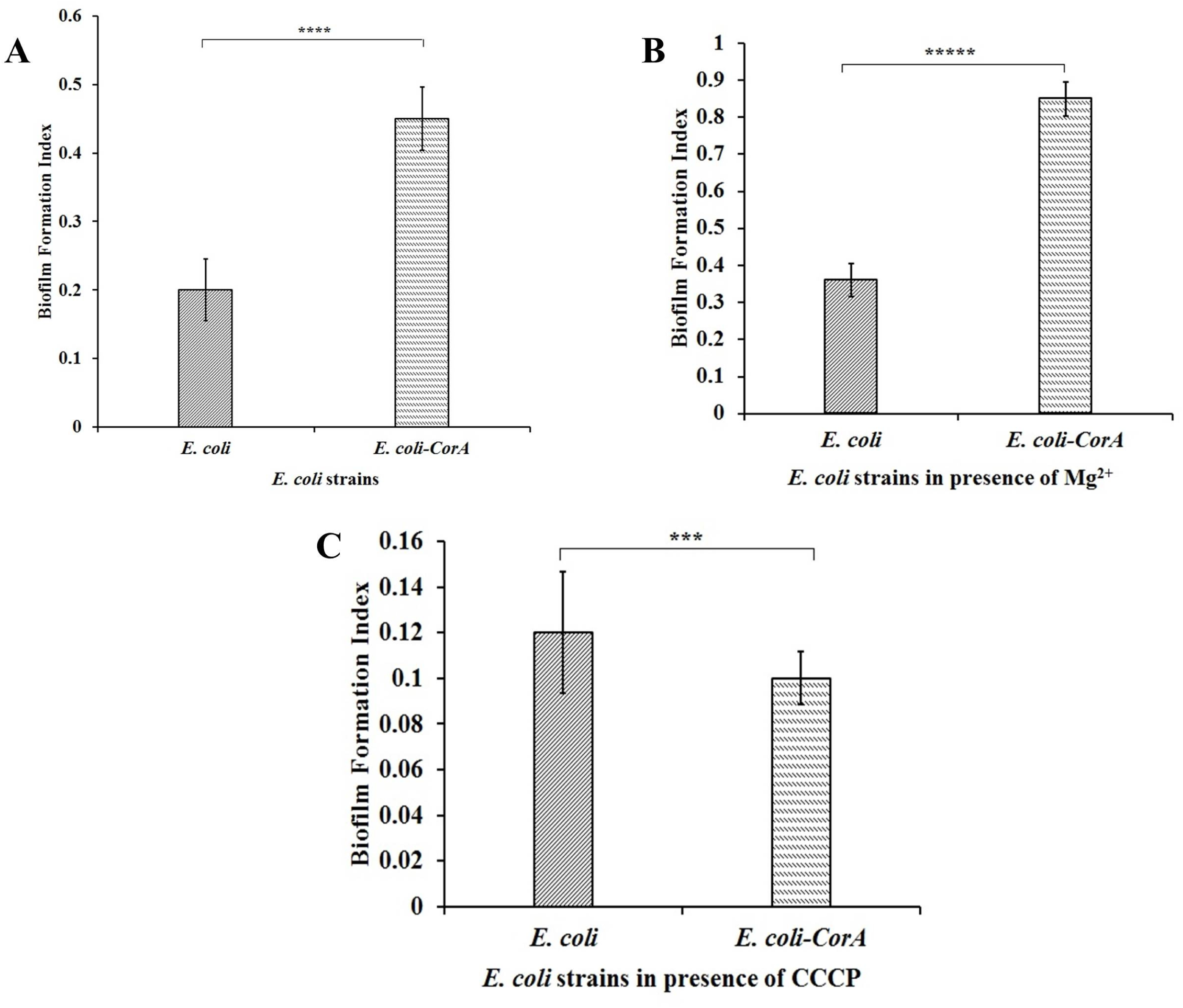
Semi-Quantitative biofilm formation assay of *E. coli*: Biofilm formation by *E. coli* CS1O9 harboring CorA (A), *E. coli* CS1O9 cells expressing CorA supplemented with Mg^2+^ (B) and CorA expressed *E. coli* cells in presence of CCCP (C) was calculated with respect to vector control by Crystal Violet staining method. The statistical significance (P value) was calculated using GraphPad by performing two samples, unpaired Student’s *t*-test where *denotes P<3;** denotes P<0.3, *** denotes P<0.03, **** denotes P<0.003, ***** denotes P<0.0003. The error bars indicate mean±SD.

**Figure 8:**
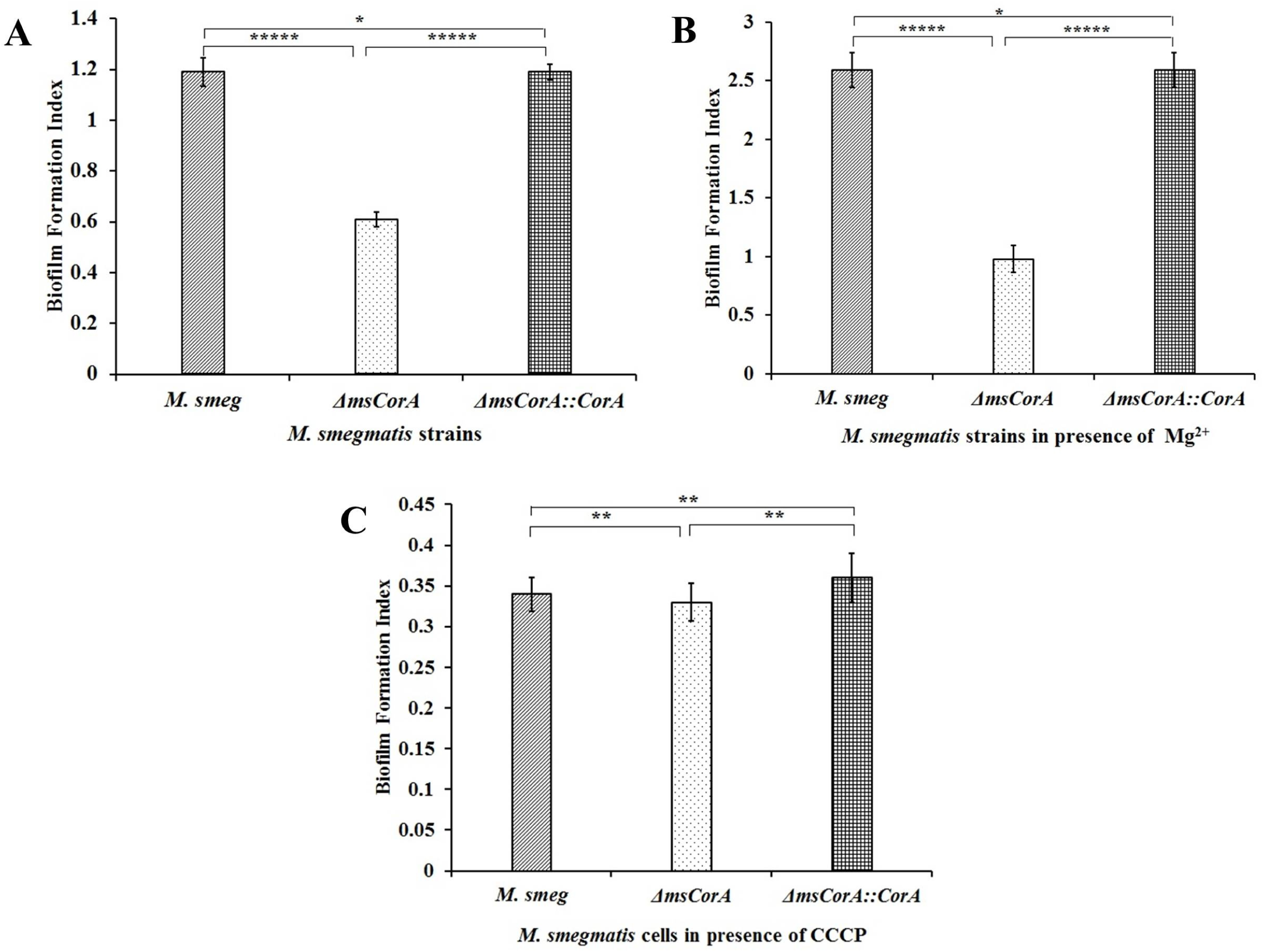
Semi-Quantitative biofilm formation assay of *M. smegmatis*: Biofilm formation by *M. smegmatis* harboring CorA (A) *M. smegmatis* cells expressing CorA supplemented with Mg^2+^ (B) and CorA expressed *M. smegmatis* cells in presence of CCCP (C) was calculated with respect to vector control by Crystal Violet staining method. The statistical significance (P value) was calculated using GraphPad by performing two samples, unpaired Student’s *t*-test where *denotes P<3;** denotes P<0.3, *** denotes P<0.03, **** denotes P<0.003, ***** denotes P<0.0003. The error bars indicate mean±SD.

**Figure 9:**
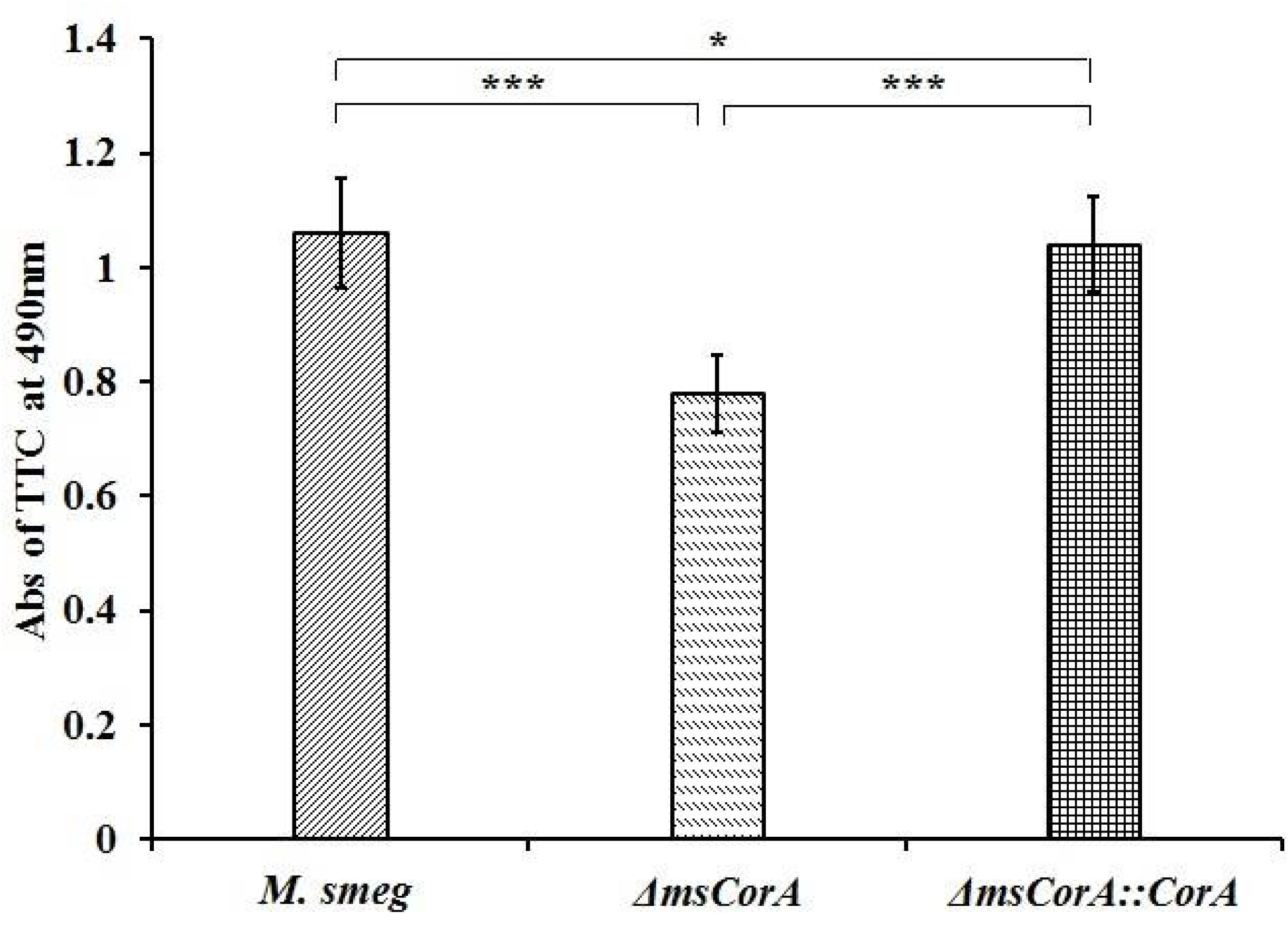
Viability of *M. smegmatis* cells in preformed biofilm: The viability of *M. smegmatis* cells in the preformed biofilm was determined by measuring the color change of the redox dye triphenyltetrazolium chloride from pale yellow to pink at 490 nm. The statistical significance (P value) was calculated using GraphPad by performing two samples, unpaired Student’s *t*-test where *denotes P<3;** denotes P<0.3, *** denotes P<0.03, **** denotes P<0.003, ***** denotes P<0.0003. The error bars indicate mean±SD.

## DISCUSSION

The feat of *Mycobacterium spp*. as an outrageous pathogen is because of its inherent capability to endure chronic infection, despite a robust host immune response (37). Macrophages of the host provide an environment for the pathogen to inhabit during infection (38). This host-pathogen interaction encourages the pathogen to undergo maturation under a hostile environment by lowering pH and limiting nutrients including magnesium (39, 40). In a living cell, Mg^2+^ is the most abundant divalent cation (41). The cells must achieve the optimum concentration of Mg^2+^ inside the cell. Magnesium transporters are employed for the said purpose and are important for various physiological processes (41). The role of magnesium in the survival of mycobacteria in host macrophages has been widely studied, though little is known about its possible role in antibiotic resistance. CorA proteins are known for magnesium uptake in bacterial species and regulating cellular magnesium concentration (42). So far, its probable role in anti-microbial and toxic compound efflux has not been widely explored. In this present work, we observed that in the presence of CorA, the susceptibilities of rifampicin and isoniazid are decreased in *M. smegmatis* and similarly, the susceptibilities of fluoroquinolones and aminoglycosides are reduced in both *M. smegmatis* and *E. coli*. This suggests that CorA has a wide range of substrate specificity varying from fluoroquinolones to anti-tubercular drugs. This finding is relevant as the above-mentioned classes of drugs are often used against tuberculosis.

Upon expression of *corA* under arabinose and tetracycline promoters led to an increase in low to moderate levels of tolerance of the host cells towards fluoroquinolones, aminoglycosides, and anti-tubercular drugs. Further to study its role in antimicrobial resistance, accumulation assays are performed. It is observed that indeed CorA influences the efflux of antibiotics and toxic dye in *E. coli* and *M. smegmatis* and CCCP plays a role as an inhibitor Intriguingly, it has been emphasized that a nickel/ cobalt efflux pump, NicT, might use the electrochemical gradient created by Ni^2+^ uptake to transport drugs, a similar observation was also noted in case of Rv3270, a Zn^2+^ transporter of *M. tuberculosis* (11, 26). Interestingly, it is observed that CorA in the presence of Mg^2+^ could enhance the tolerance of the host cells towards all the tested antibiotics varying from fluoroquinolones to aminoglycosides and anti-tubercular drugs.

Mary Ann Szegedy determined that in TM2 of CorA, conserved residues YGMNF and the hydroxyl-bearing residues Ser260, Thr270, and Ser274 are important for the metal transport through *S. typhimurium* CorA (31). Although growth phenotype generally reflected the ability to transport metal, S260 and T270 mutants had no measurable metal transport but required less Mg^2+^ to grow than MM281, the transport-deficient strain (31). The residues bearing the hydroxyl group are important for the transport of selective divalent cations and supporting optimal activity. Similarly, we could observe that mutating the similar conserved hydroxyl-bearing residues S299A and T309A of *M. smegmatis* CorA led to a loss of antimicrobial activity of CorA. It was hypothesized previously by Mary Ann Szegedy *et al*., that the mutants highly increased affinity for the metal ion, in which the cation binds too tightly for transport (31). It could be speculated that disruption in measurable metal transport may lead to the loss of Mg^2+^-facilitated resistance to drugs.

It is also noticed that the expression of *corA* also enhanced the biofilm formation of *E. coli* and *M. smegmatis*. The BFI increased by 2-4 fold for both the host cells expressing CorA. Furthermore, the *corA*-expressing cells were relatively 26% more viable than the CorA-deleted *M. smegmatis* cells (*ΔmsCorA*). Thereafter, Carbonyl cyanide 3-chlorophenylhydrazone (CCCP) an uncoupler of oxidative phosphorylation which disrupts the ionic gradient of bacterial membranes had an inhibitory effect on the biofilm formation of the host cells (43). It was perceived that CCCP not only had an inhibitory effect on the influence of multi-drug efflux by CorA but also exerted an impact on the biofilm formation of host cells negatively. Hence, it could be speculated that CorA contributes to maintaining the fitness of the microorganism, enhances the extrusion of structurally unrelated classes of antibiotics, and boosts biofilm formation of the host cells, thereby promoting antimicrobial resistance.

Based on our homology modeling and molecular docking analysis, we propose a hypothetical model for ligand transport in the CorA protein (graphical abstract). According to the literature, CorA exists in a symmetric closed conformation and multiple asymmetric open conformations. These conformations change dynamically, allowing the protein to import Mg^2+^ (44). It has also previously been studied that CorA transitions between symmetric and asymmetric states are influenced by intracellular Mg^2+^ levels. When Mg^2+^ levels are low, the closed state becomes less common, reducing the energy barrier to open states and increasing the dynamics of CorA, which facilitates the open state (45). Our *in vivo* analysis suggests that CorA might also function as an exporter of antibiotics, implying that it could act as an antiporter that imports Mg^2+^ and exports antibiotics. We speculate that in its closed symmetric conformation, antibiotics may bind to sites like C5 or C4 at the inter-subunit interfaces (**Figure S4**). When the protein transitions to different asymmetric conformations, it may allow for the export of these bound antibiotics. Docking analysis shows that most binding sites for isoniazid are located at the inter-subunit interfaces of the cytoplasmic domain, indicating that antibiotics likely move through these interfaces and enter the transmembrane pores (details are given in the supplementary section). The import of Mg^2+^ is largely governed by the conformational transitions between different asymmetric states, highlighting the importance of proper conformational transitions for Mg^2+^ import. We speculate that the same conformational transitions drive the export of antibiotics. Mutations in residues T270 and S260 in *S. typhimurium* which were previously proposed to hinder ion transport due to tight binding. In addition, cation selectivity might also restrict the transition to the open conformation, and halt Mg^2+^ transport, which may also stop antibiotic export (31). However, our model of antibiotic transport is based on preliminary analysis, and detailed structural studies considering antibiotics are required to validate this hypothesis.

## CONCLUSION

The magnesium transporter CorA of *M. smegmatis* might contribute to the development of antimicrobial resistance possibly by facilitating metal/drug cross-resistance through its efflux-pump activity.

## MATERIALS AND METHODS

### Bacterial strains, culture media, and growth conditions

The bacterial strains used in this study were *E. coli* XL1-Blue (*recA1 endA1 gyrA96 thi-1 hsdR17 supE44 relA1 lac*) (Stratagene/ Agilent, Santa Clara, CA, USA), *E. coli* CS1O9 (W1485 *glnV rpoS rph*) and *Mycobacterium smegmatis* MC^2^ 155 (ATCC). The *E. coli* strains were grown at 37 °C in Luria–Bertani broth (LB) or LB agar (Hi-Media, Mumbai, MH, India) in the presence of appropriate antibiotics (*E. coli* XL1-Blue was grown in the presence of 12 μg ml^-1^ tetracycline; strains carrying pBAD18-Cam were grown in the presence of 20 μg ml^-1^ chloramphenicol) (18). *Mycobacterium smegmatis* MC^2^ 155 strains were grown in Middle Brook 7H9 broth and 7H11 agar medium (Sigma-Aldrich, St. Louis, MO, USA) supplemented with oleic acid-ADC enrichment (Hi-Media, Mumbai, MH, India), 0.35% (w/v) glycerol, 0.05% (w/v) Tween 80 and appropriate antibiotics (50 μg ml^-1^ of kanamycin and 100 μg ml^-1^ of hygromycin for the strains carrying pMIND and pSMT-100 vectors) (19, 20). Cation-adjusted Mueller–Hinton Broth (MH) and M63 were used for antibiotic susceptibility testing of *E. coli* and *M. smegmatis* strains (HiMedia, Mumbai, MH, India), respectively. Unless otherwise specified, all the other reagents, including antibiotics, were purchased from Sigma-Aldrich (St. Louis, MO, USA).

### Construction of recombinant plasmids for *in vivo* studies

The gene CorA was amplified from the genomic DNA of *M. smegmatis* MC^2^ 155 using gene-specific primers (see **Table S1**). The PCR fragments were digested with *Nhe*I and *Hin*dIII for cloning into the pBAD18-Cam (Cam^R^) vector whereas *Bam*HI and *Pst*I were used for cloning into the pMIND (Kan^R^, Hyg^R^) vector to create the plasmid constructs pDCorA and pMCorA (18, 19). *E. coli* CS1O9 cells were transformed with pDCorA by heat-shock and *M. smegmatis* MC^2^ 155 cells were transformed with pMCorA through electroporation for the studies involving gene expression and functional characterization. Expression of the gene (CorA) cloned in pBAD18-Cam was optimized at an arabinose concentration of 0.1% (w/v) whereas expression from pMIND, a shuttle vector of *E. coli/Mycobacteria*, was optimized using 20 ng ml^-1^ of tetracycline (20).

### Genetic manipulation

The gene CorA of *M. smegmatis* was inactivated by following the method of homologous recombination discussed earlier (21). Briefly, the left and right flanking regions of CorA of *M. smegmatis* were amplified with specific primers (**Table S1**) using a thermocycler (MiniAmp Plus; Thermo Fisher Scientific, Massachusetts, United States). The amplicons were sequentially cloned in a suicide vector, pSMT-100, to create pSCorA, and electroporated in *M. smegmatis* MC^2^155 cells. The double-cross-over transformants were selected on 7H11 Agar medium supplemented with 10% OADC, 100 μg ml^-1^ of hygromycin, and 10% (w/v) sucrose. The deletion was confirmed by amplifying the pSCorA electroporated *M. smegmatis* DNA using different combinations of primers (**Table S1**) (**Figure S1**). The CorA-deleted *M. smegmatis* strain (Δ*msCorA*) was complemented with *CorA* of *M. smegmatis* cloned in tetracycline-inducible vector pMIND (Δ*msCorA*::*CorA*) and selected on kanamycin and hygromycin supplemented 7H11 agar plates for further complementation studies.

### Site-directed mutagenesis

The plasmid pDCorA was used as a template for site-directed mutagenesis. Two amino acid substitutions (S299A and T309A) were carried out using specific primers (**Table S1**) and Pfu Turbo Polymerase (Agilent Technologies, Santa Clara, CA, USA). Then amplicons were further treated with *Dpn*I (New England BioLabs, Ipswich, Massachusetts, United States). *E. coli* XL1-Blue cells were transformed with *Dpn*I-treated amplicons. All the mutations were confirmed by commercial sequencing by Eurofins Scientific (Hyderabad, TS, India). Next, CorA carrying specific mutations were cloned separately in the pMIND vector, and subsequently, the *M. smegmatis* cells were transformed with the same. Further experiments were performed to evaluate the effect of these CorA mutations on antimicrobial resistance.

### Expression of corA in E. coli and M. smegmatis

LB and 7H9 broth (10 ml each) were inoculated with 0.1% of the overnight grown culture and incubated at 37°C with a shaking speed of 150 rpm till the culture reached an OD_600_ of ∼0.2. For the expression of the gene and protein production the *E. coli* and *M. smegmatis* cultures were induced with 0.1% arabinose (w/v) and 20 ng/µL of tetracycline, respectively, and incubated for 14-16 hr. The cells were spun down at 20,000 x g for 20 mins at 4°C in Sorvall RC6 PLUS centrifuge (Thermo Scientific, Waltham, MA, USA). The supernatant was discarded and the pellet obtained was washed with 1 ml of 10mM Tris-Cl buffer (pH-7.5) and resuspended in the same buffer. The protease inhibitor, phenylmethylsulfonyl fluoride (PMSF), was added at a final concentration of 1 mM to the cell suspension, and it was sonicated with five pulses of 45 sec each for *E. coli* and 60 sec for *M. smegmatis*, respectively, in a Corning CoolRack M6 placed in ice, followed by centrifugation at 20,000 x g for 5 mins. The supernatant fraction was collected and further centrifuged at 4°C for 1 hr. at 20,000 x g. The supernatant was separated and the pellet fraction was further resuspended in 100 µL of Tris-Cl buffer (pH-7.5). The pellet fraction was treated with 2% sarcosyl (sodium lauroyl sarcosinate) (w/v), mixed thoroughly, and incubated at 37°C on a thermomixer (Eppendorf, Hamburg, Germany) with shaking for 1 hr. to solubilize the membrane proteins. The sample was further centrifuged 20,000 x g for 1 hr. at 4°C and the supernatant was collected. The proteins in the collected supernatant were estimated, and analyzed by running through sodium dodecyl sulfate polyacrylamide gel electrophoresis (SDS-PAGE) (12% acrylamide) (20, 22).

### Determination of susceptibility towards different antibiotics, dyes, and metal salts

Susceptibilities of the strains towards different antibiotics, *viz*., fluoroquinolones, aminoglycosides, and anti-tubercular drugs, dyes such as Ethidium Bromide (EtBr), Nile red, acriflavine, rhodamine B, and metal salts of magnesium, zinc and cobalt, were determined using micro-broth dilution method, and assessed according to Clinical and Laboratory Standards Institute (CLSI) guideline. The assays were performed in 96-well micro-titre plates with an assay volume of 300µl per well for *E. coli* strains, and 200µl per well for *M. smegmatis* strains (11). Two-fold serial dilutions of different antibiotics were made in cation-adjusted MH broth and 7H9 media where each well was inoculated with 10^5^ cells. The plates were incubated at 37 °C for 16–18h for *E. coli* and 48–72h for *M. smegmatis*, respectively. MgCl_2_ was added at 1/8th of its MIC, and 100 µM of CCCP was added, to the MH broth and 7H9 media, respectively, to study its effect on antimicrobial susceptibility. Bacterial growth was assessed at *A*_600nm_ by Multiskan Spectrum spectrophotometer (model 1500; Thermo Scientific, Waltham, MA, USA). The assays were performed in triplicates and repeated six times to report consistent results.

### Ethidium Bromide accumulation assay

The *M. smegmatis* cells (10^5^ cells) were washed with phosphate buffer (PB buffer) supplemented with 0.05% Tween 80 and MgCl_2._ The optical density of the suspension was adjusted at *A*_600nm_ to 0.4 with PB buffer, and glucose was added at a final concentration of 0.4% The cultures (95 µL) were distributed in 96-well micro-titre plates and EtBr was added at a final concentration of 1.5 µM. Relative fluorescence was measured by BioTek Cytation 5 Cell Imaging Multimode Reader (Agilent Technologies, Santa Clara, CA, USA) with an excitation wavelength of 530 nm and emission wavelength of 560-640 nm. Fluorescence data were acquired every 60 seconds for 15 minutes at 37°C (23).

### Determination of Intracellular antibiotic accumulation activity

Intracellular accumulation of norfloxacin and ofloxacin (fluoroquinolones antibiotics), were determined as described previously (11). *E. coli* and *M. smegmatis* were grown to 10^5^ cells after inducing with 0.1% arabinose and 20 ng/µL tetracycline, respectively. The cells were washed thrice with 50 mM PB (pH 7.2). *E. coli* cells were energized with 0.2% (w/v) glucose for 20 min at 30 °C. The antibiotics were added at a final concentration of 10 mg L^-1^ to the cell suspensions and aliquots (0.5 ml) were drawn at intervals of 5 min. To check for the energy dependency in the efflux process, CCCP (100µM) was added to the cell suspensions after 15 min, and aliquots were drawn further. MgCl_2_ was added at a concentration of 1/8^th^ of its MIC value for the assessment of its effect on the efflux activity of cells expressing *corA.* Cells were harvested immediately after aliquoting, washed thrice with PB, and resuspended in 0.1 M glycine hydrochloride, pH-3.0. Subsequently, *E. coli* and *M. smegmatis* cells were incubated at 37 °C for 60 min and overnight, respectively (24). The suspensions were spun down, and the fluorescence of the supernatant was measured by a spectrofluorimeter (FluoroMax 4, HORIBA Scientific Instruments, Irvine CA, USA) at an excitation wavelength of 281 nm, emission wavelength 447 nm for norfloxacin, excitation wavelength of 280 nm and emission wavelength of 485 nm for ofloxacin, respectively. The relative norfloxacin efflux of the wild-type and mutant strains were calculated, which is given as follows: RE (relative efflux) = 1+ (N_control_ − N_test_)/N_control_. In this equation, N_control_ represents the antibiotic uptake of vector control and N_test_ represents antibiotic uptake by test strains (25). The relative percentage of efflux was calculated by: (N_control_ - N_test_)/ N_control_ X 100 (26).

### Growth curve analysis

Overnight cultures were washed with PB buffer and diluted to *A*_600nm_ equivalent to McFarland number 0.5 and inoculated to 100 ml 7H9 broth supplemented with 0.05% Tween 80, and 100 μg ml^-1^ hygromycin 50 μg ml^-1^ kanamycin. The cultures were grown at 37°C, and the CFU per milliliter (CFU/ml) was calculated from 0 to 40 h at a growth interval of 3 h. The generation time was calculated for each sample from the linear part of the log phase in the growth curve (20, 26).

### Semi-quantitative crystal Violet-based biofilm assay

Biofilm formation was assessed by using the static 24-well plate microtitre-based biofilm assay (24, 25, 27). The wells of a sterile 24-well polystyrene microtitre plate were filled with 1 ml 1/5 LB and 1XM63 medium that was inoculated with 10^5^ cells, and subsequently, incubated at 37 °C for 48 hr. for biofilm formation under static conditions. Following this, the wells were washed thrice with Phosphate Buffer Saline (PBS) to remove the planktonic cells and stained with 0.1 % crystal violet (CV) stain for 15 min. Following staining the excess CV was removed by washing with water. The bound dye was solubilized in 1 ml of 33 % acetic acid (v/v), and the OD of the acetic acid-dye solution was measured at 600 nm using an ultraviolet/ visible spectrophotometer (Multiskan Spectrum-1500 Spectrophotometer [Thermo Scientific, Switzerland]). The biofilm formation index (BFI) was calculated by the formula: BFI= (OD_600_ of the acetic acid solubilized CV of culture – OD_600_ of the acetic acid solubilized CV of control) / OD_600_ of planktonic cells). Further, to determine the effect of CCCP and magnesium on the biofilm-forming ability of the host strains, the wells of the 24-well plate were filled with 1 ml of M63 medium, supplemented with CCCP and 1/8^th^ MIC of Mg^2+^, and inoculated with *M. smegmatis* cells. The plates were incubated for 3 days at 37°C. After incubation, the wells were washed with sterile PBS and stained with 0.1% crystal violet dye for 15 mins. The biofilms formed in the wells were washed again and destained with 33% glacial acetic acid. The absorbance was measured at OD_600_ using a Multiskan Spectrum. Biofilm formation index was calculated to determine the effect of Mg^2+^ by BFI= (OD_600_ of the acetic acid solubilized CV of culture – OD_600_ of the acetic acid solubilized CV of control) / OD_600_ of planktonic cells supplemented with magnesium. The percentage of biofilm inhibition was calculated = [A_control_- A_treated_)/A_control_] X 100 (26). Additionally, to ascertain the relative difference in the amount of viable cells contributing to biofilm formed, the wells of 24 well plates were filled with M63 medium and inoculated with 10^5^ cells of *M. smegmatis*. The plate was incubated for 3 days at 37°C. After incubation, the wells were washed with sterile PBS, spent media was discarded, the wells were refilled with M63 medium and incubated further for 36 hr. The viability of the cells was detected with the help of the redox dye triphenyl tetrazolium chloride (TTC) used at a final concentration of 0.05%. In the presence of viable cells, the dye turns from pale yellow to pink. The color was measured using a Multiskan Spectrum spectrophotometer at A_490nm._ The percentage of dead cells was determined via [A_control_-A_treated_)/A_control_] X 100 (28).

### Phase-contrast microscopic analysis of biofilm formation of *M. smegmatis* cells

The wells of 24 well plate polystyrene plates fitted with microscopic cover glass slips were filled with 1XM63 medium inoculated with 10^5^ cells and incubated at 37°C for 3 days in static condition. The wells were washed with PBS buffer and stained with 0.1% crystal violet solution for 15 min and finally, the wells were washed with water. The biofilms formed on the coverslips were subsequently visualized using an OLYMPUS 1×71 (Olympus Corporation, Tokyo, Japan) (29).

### Statistical Analysis

All experiments related to the accumulation of antibiotics and dyes were performed in triplicate and the results were calculated as mean±standard deviation. The statistical significance (P value) was calculated using GraphPad by performing two samples, unpaired Student’s *t*-test where *denotes P<3;** denotes P<0.3, *** denotes P<0.03, **** denotes P<0.003, ***** denotes P<0.0003.

## ACKNOWLEDGEMENT

We thank Dr. Somdeb Bose Dasgupta, Bioscience and Biotechnology, IIT Kharagpur, India for the BioTek Cytation 5 Cell Imaging Multimode Reader (Agilent Technologies, Santa Clara, CA, USA) facility.

## RESEARCH CONTRIBUTION

D.C. and A.S.G designed the experiments and analyzed the results. D.C. performed most of the experiments and wrote the draft of the manuscript, A.S.G was involved in the overall designing and supervision of the work and editing of the manuscript, A.R.D.M was involved in performing *E. coli* related experiments, A.P.P did the *in silico* experiments and analysis, S.K.R was involved in initial designing of the project and data analysis; B.B did the biofilm microscopy analysis, and D.B helped in data analysis.

